# Multiple approaches for CRISPR-based targeting of DNA methylation to promoters of bacterial and viral susceptibility genes in cassava

**DOI:** 10.64898/2026.06.04.730177

**Authors:** Kerrigan B. Gilbert, Zuh-Jyh Daniel Lin, Kira M. Veley, Myia K. Stanton, Marisa Yoder, Joanna Norton, Suhua Feng, Yan He, Gabriela L. Hernandez, Greg Jensen, Emma Wozniak, Ke Ke, Sharon Motomura-Wages, Steven E. Jacobsen, Rebecca S. Bart, James C. Carrington

## Abstract

Targeted epigenetic modifications of specific gene regulatory regions have the potential to confer beneficial traits for crop improvement. Two recently developed CRISPR/Cas9-based epigenome editing tools were tested in transgenic cassava to target cytosine methylation to the promoter region of *MeSWEET10a,* a necessary gene for infection by the Cassava bacterial blight pathogen, *Xanthomonas phaseoli* pv. *manihotis*. The two systems leverage unique methyltransferases, and each induced distinct DNA methylation profiles at the targeted site, decreased effector-triggered *MeSWEET10a* expression, and attenuated water-soaking symptoms in inoculated leaves. Further, DNA methylation was simultaneously targeted, in addition to *MeSWEET10a*, to two susceptibility genes for Cassava brown streak virus. Relative levels of *de novo* DNA methylation at the three loci were inversely correlated with DNA methylation-antagonizing H3K4me3 marks. Finally, an initial assessment of DNA methylation after one generation indicated specific inheritance of CpG methylation that was unstable in the absence of the methyltransferase systems.

## Introduction

Cassava (*Manihot esculenta*) is a critical dietary staple for an estimated 800 million people in the tropics (Food and Agriculture Organization of the United Nations 2018). In sub-Saharan Africa, cassava production is significantly affected by cassava mosaic and cassava brown streak diseases (CMD & CBSD), caused by cassava mosaic geminiviruses and cassava brown streak ipomoviruses, respectively (Food and Agriculture Organization of the United Nations 2018; Legg et al. 2015). Cassava varieties with high levels of resistance to CMD are available, while CBSD resistant cassava varieties are under development (Legg et al. 2015; Wagaba et al. 2016; Beyene et al. 2016). Development and deployment of CBSD resistant cassava is particularly crucial as the disease continues to spread through and towards the first and second-most productive cassava growing countries, the Democratic Republic of Congo and Nigeria, respectively (FAO 2023; Chikoti and Tembo 2022). A bacterial disease, cassava bacterial blight (CBB), caused by *Xanthomonas phaseoli* pv. *manihotis* (Xpm; formerly *X. axonopodis* pv. *manihotis* (Xam)), is also an important constraint on global cassava production. The incidence of CBB is sporadic and unpredictable, however, affected cassava fields can suffer up to 100% loss (Lin et al. 2019; López and Bernal 2012; Yaméogo et al. 2024). Cassava varieties with varying levels of strain-specific resistance to Xpm exist but have not been introduced into Africa-adapted varieties (López and Bernal 2012; Restrepo et al. 2004).

Susceptibility to pathogens is mediated by their manipulation of hosts at the molecular level. While plants have evolved active modes of resistance against pathogens, such as through immune receptor-mediated responses, resistance can also be conferred by disrupting the ability of pathogens to hijack host processes needed for progression of disease. Host genes encoding proteins that are hijacked by pathogens for disease are known as susceptibility (S) genes (Baruah et al. 2025; Garcia-Ruiz et al. 2021). Both Xpm and cassava brown streak ipomoviruses require S genes for infection and disease. Like many bacterial plant pathogens, Xpm uses the type III secretion system to deliver effector proteins into the plant apoplast for various virulence functions. A subset of these are xanthomonad specific transcription activator-like effectors (TAL Effectors) that bind to the promoters of specific S genes and induce their expression. In particular, activation of the S gene *MeSWEET10a* by the TAL effector TAL20 is critical. Frameshift mutations in *MeSWEET10a* coding sequence, or mutations affecting the TAL20 effector binding element (EBE) upstream of *MeSWEET10a,* strongly attenuate water-soaking associated with CBB infection (Elliott et al. 2024). Viruses belonging to the family *Potyviridae*, like cassava brown streak ipomovirus (CBSV), often coopt specific host-encoded eIF4E translation initiation factors through interaction with the virus-encoded genome-linked protein (VPg); mutants with defects in relevant eIF4E family members may lose susceptibility to infection (Robaglia and Caranta 2006; Bastet et al. 2017). For example, simultaneously knocking out the *eIF4E*-family genes *nCBP-1* and *nCBP-2* in cassava strongly reduces cassava brown streak virus accumulation and necrosis in storage roots (Gomez et al. 2019; Z. J. D. Lin et al. 2025).

These examples of S gene modification resulting in resistance were enabled by the advent of CRISPR/Cas genome editing techniques. CRISPR/Cas technologies have accelerated functional genetic studies and translation of fundamental disease resistance principles into orphan crops like cassava. In addition to S gene editing for resistance, epigenome editing is emerging as a potential strategy for modulating S gene expression (Veley et al. 2023). Repressive epigenetic marks, both in the form of promoter cytosine methylation and certain associated histone post-translational modifications, can now be established *de novo* in plants by targeting DNA methyltransferases and histone modifying enzymes to promoters of interest (Papikian et al. 2019; Ghoshal et al. 2021; Wang et al. 2025). These tools have been extensively tested at the *FWA* locus in *Arabidopsis* and have been demonstrated to restore heritable epigenetic repression of *FWA* that has been lost in the *fwa-4* epimutant. In terms of controlling xanthomonad diseases, DNA methylation of TAL effector EBEs alone is sufficient for disrupting TAL effector binding and preventing activation of S genes, which we previously demonstrated with Xpm and *MeSWEET10a* in cassava (Veley et al. 2023; Deng et al. 2012). This was done by designing an artificial zinc finger that binds the TAL20 EBE and fusing it to DEFECTIVE IN MERISTEM SILENCING 3 (DMS3), a component of the RNA-directed DNA methylation pathway. The knockdown effect of promoter epigenetic editing may be advantageous in cases where S genes redundantly contribute to disease and cannot be knocked out due to being essential for plant viability, as evidence suggests for the *eIF4E*-family of genes in cassava (Gomez et al. 2019; Z. J. D. Lin et al. 2025).

Here we describe the application of two recent iterations of guide RNA-directed epigenome editing tools in cassava for resistance to CBB and CBSD. Both harness a deactivated Cas9 enzyme and two gRNAs specific to cassava per target locus. The first construct fused the dCas9 protein to ten copies of the SunTag epitope system which in turn recruits the catalytic domain of the methyltransferase DRM (DOMAINS REARRANGED METHYLTRANSFERASE) from *Nicotiana benthamiana* (Papikian et al. 2019); this construct is referred to as dCas9-SunTag-DRMcd. The second technology has a variant of the CG-specific methyltransferase MQ1 fused directly to dCas9; this construct is referred to as dCas9-MQ1v (Ghoshal et al. 2021).

## Results

### DRMcd-mediated *de novo* methylation at *MeSWEET10a* promoter and attenuation of CBB symptoms

Previously we used an engineered ZF fused to DMS3 to target de novo methylation to the *MeSWEET10a* promoter in cassava, which includes the TAL20 binding site of Xpm. The catalytic domain of DRM2, as part of a deactivated Cas9 (dCas9) SunTag system, has been effective in establishing methylation at the *FWA* promoter in the Arabidopsis *fwa4* epimutant <u>(Papikian et al. 2019)</u>. As such, transgenic cassava lines were generated expressing dCas9-SunTag with either DRMcd or without an enzyme reagent (ΔDRM) (Figure 1a). The complex was targeted to the *MeSWEET10a* promoter using two gRNAs that flank the Effector Binding Element (EBE) sequence (Figure 1b). Three independent transformed lines were isolated expressing DRMcd, each containing only one T-DNA insertion (Supplemental Figure 1a-d), and one transformed line expressing the ΔDRM negative control.

**Figure 1:**
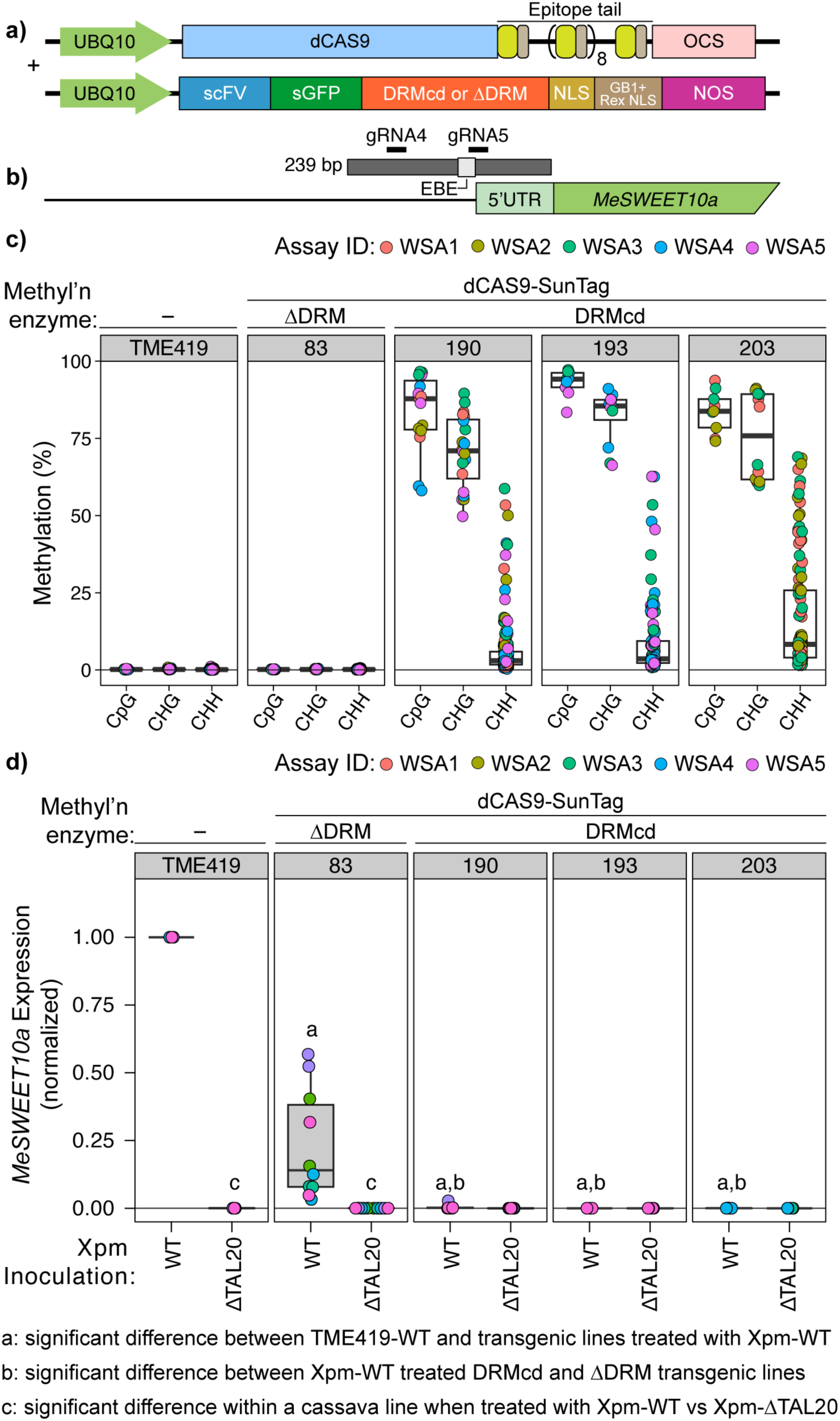
The methyltransferase DRM targets *de novo* methylation to the cassava *MeSWEET10a* promoter and reduces induction of *MeSWEET10a* during infection by Xpm. (a) Schematic of the dCAS9-SunTag system + DRMcd cassettes; modified from Papikian et al. (2019); definition of abbreviations are at the end of the figure legend. (b) Schematic of the *MeSWEET10a* (Manes.06G123400) promoter region; the 239 bp area in dark grey contains the two gRNAs used to target dCAS9 (gRNA4 and gRNA5) and the effector binding element (EBE) recognized by TAL20. (c and d) Molecular characterization of epi-edited cassava lines. (c) Methylation signal within the 239 bp region of the *MeSWEET10a* promoter shown in (b), measured by PCR-based bisulfite sequencing (ampBS-seq). Data from five independent water soaking assay (WSA) experiments is plotted and color-coded by Assay ID. Each dot represents the average methylation signal at a given location within the region of interest from two leaves from the same plant per sample per assay. Cytosine methylation was measured in the three contexts: CpG, CHG, and CHH. (d) Induction of *MeSWEET10a* expression following infiltration by either wildtype Xpm (WT) or Xpm without the TAL20 effector (ΔTAL20). Expression was evaluated using qPCR and the cassava genes GTPb (Manes.09G086600) and PP2A4 (Manes.09G039900) were used as internal controls. Two technical replicates were measured and plotted for each individual plant per sample per assay, with data from up to five independent assays present. Results from two-sided Welch’s t-test *p* values are noted and comparisons are defined at the bottom of the figure. UBQ10: Arabidopsis UBIQUITIN10 promoter; OCS: octopine synthase terminator; scFv: single chain variable fragment; sGFP: superfolder-GFP; DRMcd: catalytic domain of DOMAINS REARRANGED METHYLTRANSFERASE 2 (DRM); NLS: nuclear localization signal; GB1: solubility tag; Rex NSL: nuclear localization signal; NOS: Nopaline synthase terminator.

Amplicon-based bisulfite sequencing (ampBS-Seq) of the *MeSWEET10a* promoter region of wild-type and transgenic plants showed that lines expressing the DRMcd domain had robust *de novo* methylation while no methylation was observed in either the wild type or negative control line (Figure 1c). In particular, methylation levels at the three CpG residues within the region were generally over 90% in each independent line while CHG and CHH methylation was also present.

As binding of the TAL20 effector is known to induce upregulation of *MeSWEET10a*, we evaluated individual leaves from each cassava line for *MeSWEET10a* expression following infection by Xpm668 to determine if the presence of methylation correlated with reduced induction. Compared to the wild-type plants, all transgenic lines had significantly lower induction of *MeSWEET10a* (Figure 1d), including the unmethylated ΔDRM control line. This observation could be explained by CRISPR interference (CRISPRi), if binding of the dCas9 protein to the *MeSWEET10a* promoter region physically blocks the TAL20 effector from effectively accessing the EBE binding site; indeed, CRISPRi has been previously described in the ΔDRM line 82 (Z.-J. D. Lin et al. 2025). Effects on the induction of *MeSWEET10a* appeared variable across the replicates of the ΔDRM line, indicating the presumed effect of CRISPRi is inconsistent. CRISPRi was also likely acting in the methylated, DRMcd transgenic lines, however a more consistent and stronger lack of induction of *MeSWEET10a* expression was observed in all methylated individuals, with induction at or near 0 in multiple replicates for all three lines.

These cassava lines were also subjected to quantitative image analysis of water-soaking lesions caused by infection with either wild-type Xpm668 or the ΔTAL20 mutant; a representative image of one of the lesions for each treatment is shown in Figure 2a. Lesions were measured for two different characteristics: total area and grey-scale color intensity (Elliott et al. 2024). The lesion area for plants with methylation at the *MeSWEET10a* promoter were significantly smaller than the wild type and unmethylated transgenic control (Figure 2b). Further, as expected, infections by the Xpm ΔTAL20 mutant caused significantly smaller lesions as compared to wild-type Xpm in only the wild-type and ΔDRM plants; the difference in size within the methylated lines was not significant. Similarly, evaluation of grey-scale intensity of the lesions revealed that wild-type and unmethylated plants had significantly darker lesions than individuals with methylation (Figure 2c) while the intensity of the lesions in methylated plants was generally not significantly between wild-type and ΔTAL20 inoculations. Despite the significant impact on *MeSWEET10a* induction in the unmethylated ΔDRM control, lesion area and greyscale intensity were not significantly different than wild type.

**Figure 2:**
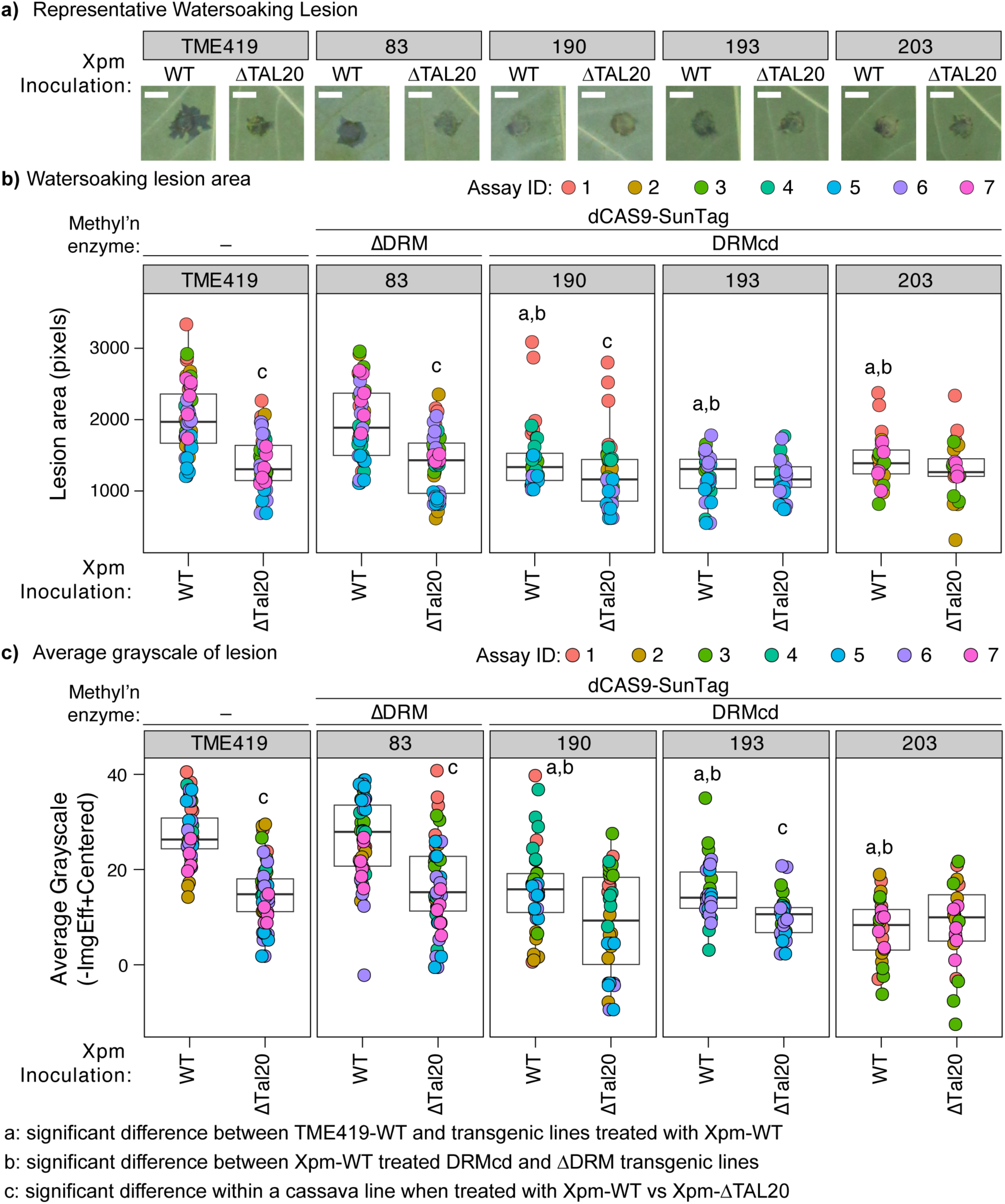
DRMcd-methylated cassava lines have attenuated CBB symptoms. (a) Representative images of water-soaking lesions on leaves from TME419 WT, dCAS9-SunTag-DRMcd, and dCAS9-SunTag-ΔDRM expressing plants. Images were taken 6 days post-infection by either Xpm668 (WT) or a Xpm668 TAL20 deletion mutant (ΔTAL20). Scale bar = 0.5 cm. (b and c) Evaluation of water-soaking lesions following infection by either Xpm668 (WT) or Xpm668 TAL20 deletion mutant (ΔTAL20). Three leaves per plant were inoculated and the resulting lesion evaluated across up to seven assays. (b) Area of lesion (in pixels) and (c) intensity of water-soaking lesions (-negative mean gray-scale value for the water-soaked region relative to the average of the mock-treated samples). Calculated *p* values from two-sided Kolmogorov–Smirnov test are shown on (b and c) and are defined at the bottom of the figure.

Together these results demonstrate that the dCas9-SunTag-DRMcd complex was able to successfully introduce *de novo* methylation in multiple, independent transgenic cassava lines, and that only in the presence of methylation was activation of *MeSWEET10a* suppressed with a corresponding reduction in CBB-related symptoms.

### MQ1v-mediated CpG methylation at *MeSWEET10a* promoter and attenuation of CBB symptoms

CG methylation is thought to be efficiently propagated during cell division (Ghoshal et al. 2021). As such, we tested the ability of a CG-specific methyltransferase fused to dCas9 to establish CG-specific, *de novo* methylation at the TAL20 EBE region and affect Xpm-associated phenotypes. Transgenic cassava plants were generated containing either dCas9 directly fused to a variant of the bacterial CpG methyltransferase MQ1 (MQ1v) or to a deactivated version of MQ1 (dMQ1) (Figure 3a). The fused proteins were targeted to the *MeSWEET10a* promoter using the same gRNAs as the dCas9-SunTag-DRMcd construct (Figure 1b). Three independent transformed lines were isolated expressing dCas9-MQ1v (line IDs 5, 22, and 76) and two expressing the inactive enzyme dCas9-dMQ1 (line IDs 122 and 127). Each dCas9-MQ1v line contains a single copy of the T-DNA insertion (Supplemental Figure 1e).

**Figure 3:**
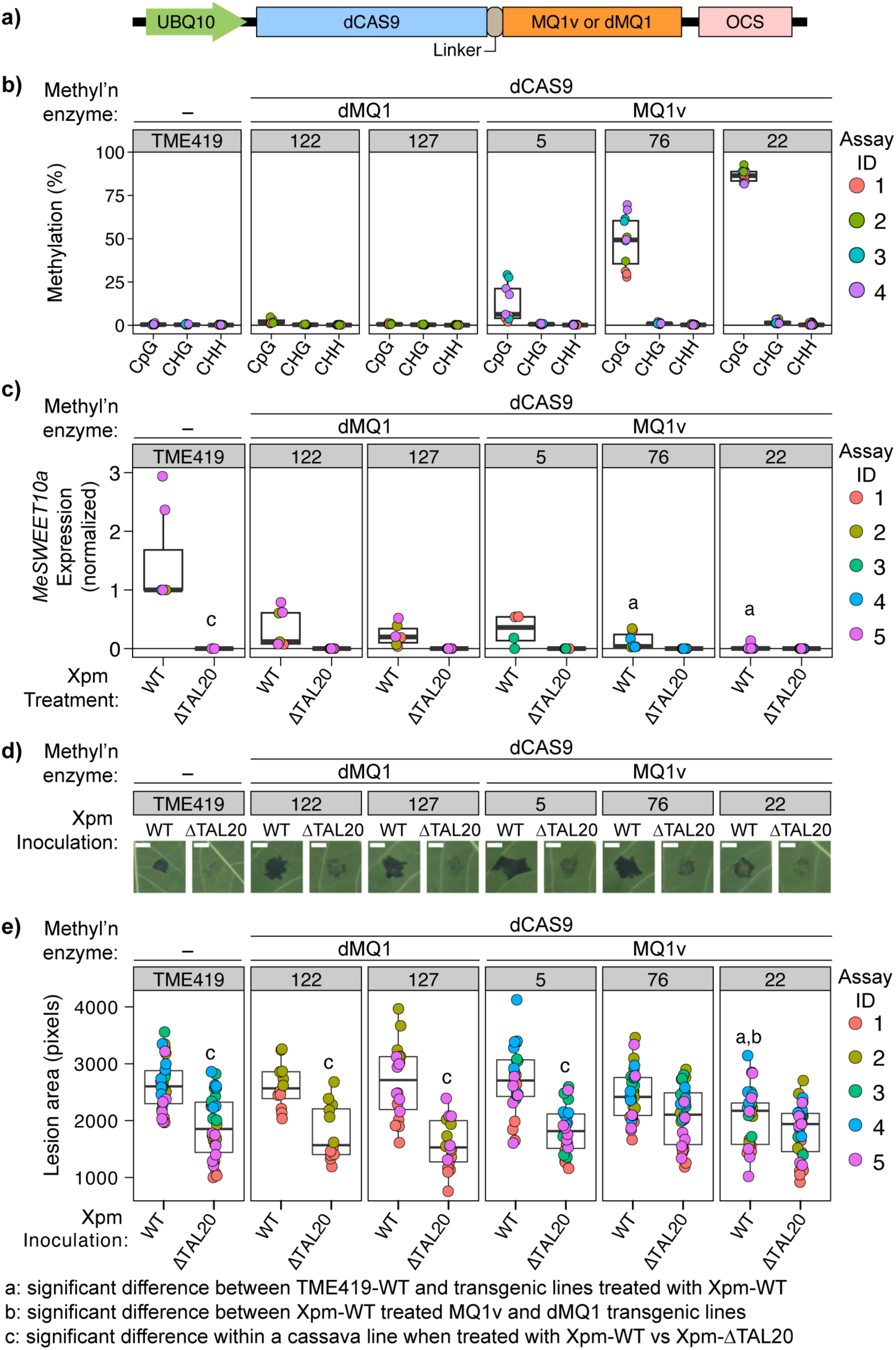
The methyltransferase MQ1v targets CG methylation to the cassava *MeSWEET10a* promoter and attenuates CBB symptoms. (a) Schematic diagram of the dCas9-based construct, containing either MQ1v or the deactivated MQ1 (dMQ1) enzyme. (b) Methylation within the *MeSWEET10a* promoter region measured by PCR-based bisulfite sequencing (ampBS-seq). Data from four independent experiments is plotted, color-coded by Assay ID. Each dot represents the average methylation signal at a given location within the region of interest from two leaves from the same plant per sample per assay. Cytosine methylation was measured in the three contexts: CpG, CHG, and CHH. (c) Induction of *MeSWEET10a* expression following infiltration by either wildtype Xpm (WT) or Xpm without the TAL20 effector (ΔTAL20). Expression was evaluated using qPCR and the cassava genes GTPb (Manes.09G086600) and PP2A4 (Manes.09G039900) were used as internal controls. Two technical replicates were measured and plotted for each individual plant per sample per assay, with data from at least four independent assays for each sample. Results from two-sided Welch’s t-test *p* values are indicated and the definitions are indicated at the bottom of the figure. (d) Representative images of water-soaking lesions on leaves from TME419 WT, dCAS9-dMQ1, and dCAS9-MQ1v expressing plants. Images were taken 6 days post-infection by either Xpm668 (WT) or a Xpm668 TAL20 deletion mutant (ΔTAL20). Scale bar = 0.5 cm. (e) Area of the water-soaking lesions following infection by either Xpm668 (WT) or Xpm668 TAL20 deletion mutant (ΔTAL20). Data from three lesions per leaf per assay are plotted. Calculated *p* values from two-sided Kolmogorov–Smirnov test are shown and defined at the bottom of the figure.

Transgenic lines expressing the active methyltransferase showed *de novo* methylation, while both the wild-type and deactivated MQ1 lines did not, across four different assays (Figure 3b). Specifically, only CpG methylation was observed in the three dCas9-MQ1v lines, although the level of methylation was variable, ranging from below 10% to nearly 90%. Quantitative RT-PCR evaluated the impact of the *de novo* methylation to hinder induction of *MeSWEET10a* during infection by both wild-type and the ΔTAL20 mutant Xpm. The level of *MeSWEET10a* expression was roughly inversely correlated with methylation, where the line with the highest methylation had the lowest induction and the line with the lowest methylation had the highest induction of *MeSWEET10a* (Figure 3c). Similar to the ΔDRM control line, a reduction in gene induction was observed in the two deactivated MQ1 lines suggesting CRISPRi by the dCas9-dMQ1 complex, however these were not statistically different from the induction observed in Xpm-treated TME419. Two of the three lines with *de novo* methylation had significantly reduced induction of *MeSWEET10a* following inoculation by wild-type Xpm, specifically the line with the highest and intermediate levels of methylation, suggesting that moderate levels of methylation may negatively impact *MeSWEET10a* expression, particularly in combination with CRISPRi.

Water soaking phenotype after Xpm infections were evaluated by quantitative image-based analyses. Representative lesions from each treatment are shown in Figure 3d. Only leaves from Line 22, with the highest levels of *de novo* methylation within the *MeSWEET10a* promoter, had significantly reduced lesion area and lesion gray-scale intensity in comparison to the controls (Figure 3f and Supplemental Figure 2). Neither of the other two lines expressing MQ1v nor the unmethylated dMQ1 transgenic lines showed a significant decrease in either area or color intensity in comparison to wild-type values. Together these results demonstrate that strong methylation in only the CpG context is sufficient to result in significant improvement with respect to CBB symptoms. Specifically, attenuated symptoms are observed when as few as three cytosines in the CpG context are methylated.

### Differences in *de novo* methylation at the *MeSWEET10a* and *nCBP* promoters simultaneously targeted using DRMcd

The cassava *nCBP*-*1* and *nCBP-2* genes contribute redundantly to susceptibility of CBSV and development of necrotic storage root symptoms in cassava. Mutations in these genes result in increased resistance to CBSV (Gomez et al. 2019; Z. J. D. Lin et al. 2025). These two genes, and the *MeSWEET10a* promoter, were simultaneously targeted for *de novo* methylation using the transgenic dCas9-SunTag-DRMcd system (Figure 4a and 4b), resulting in two transgenic lines. Only modest levels of *de novo* methylation in CHH and CHG contexts were detected at *nCBP-1* and *nCBP-2* loci, with the maximum percentage of methylated cytosines at any given nucleotide position peaking at approximately 80% while negligible levels of CpG methylation were detected (Figure 4c). In contrast, robust *de novo* methylation was again observed at the *MeSWEET10a* locus, including strong CpG methylation signals (Figure 4c). Similarly, CHG methylation was consistently higher at the *MeSWEET10a* locus than either of the *nCBP* genes (Supplemental Figure 3). The differences between methylation at the *nCBP-1*, *nCBP-2*, and MeSWEET10a loci were consistent across both independent transgenic individuals, as well as after multiple clonal propagations.

**Figure 4:**
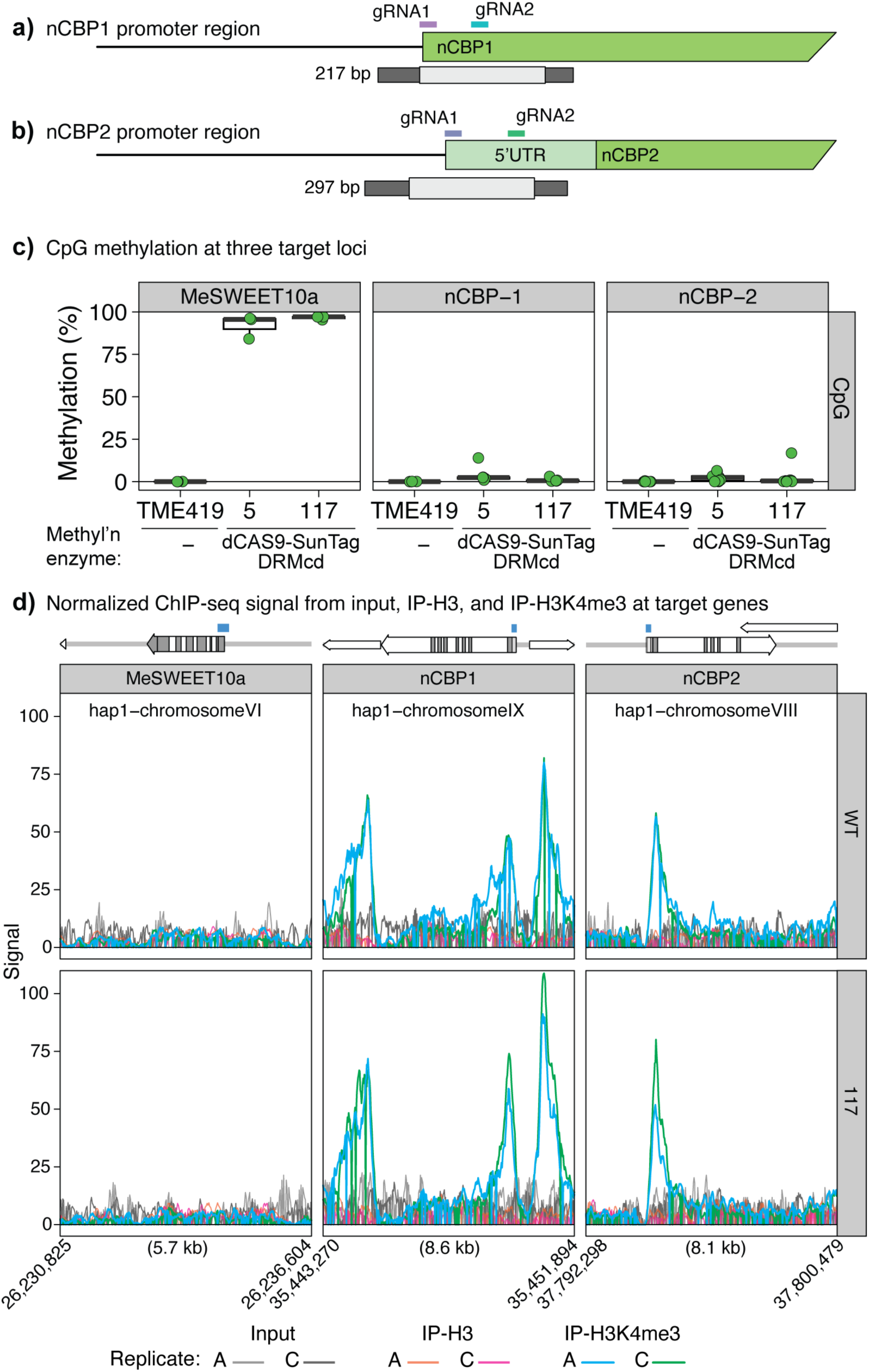
Simultaneous targeting of *de novo* methylation to three promoters in cassava using the dCAS9-SunTag-DRMcd system. (a) Schematic of the *nCBP1* promoter region (Manes.09G140300) and (b) schematic of the *nCBP2* promoter region (Manes.08G145200) targeted for methylation; gene structure based on the version 6 genome of *M. esculenta* at Phytozome. *MeSWEET10a* targeting reagents the same as in Figure 1a. Dark grey rectangle denotes the PCR amplicon used in ampBS-seq; light grey rectangle denotes the region plotted in (c) and Supplemental Figure 3. Location of gRNAs, 5’UTR, and first exon of the genes is labelled. (c) Methylation at the *MeSWEET10a*, *nCBP-1*, and *nCBP-2* promoter regions in the CpG context as measured by PCR-based bisulfite sequencing (ampBS-seq) in two independent transgenic lines expressing dCas9-SunTag-DRMcd (5 and 117) and wild type TME419. Y-axis is percent methylated from 0 to 100. (d) Normalized H3K4me3, H3, and input signal from ChIP-seq at the *MeSWEET10a*, *nCBP-1*, and *nCBP-2* regions. Gene models for the three genes of interest with exons in dark grey are along the top; blue boxes are the same region as plotted in (c); nearby genes are shown without exons. Two ChIP-seq replicates, A and C, are plotted for both wild type TME419 and dCas9-SunTag-DRMcd line 117.

### Histone marks that antagonize DNA methylation are enriched at the *nCBP-1 and nCBP-2* promoters

The consistent differences between targeted methylation at the *nCBP* promoters and *MeSWEET10a* promoter could be due to several possible reasons, including differences in antagonistic or permissive features of chromatin at the targeted sites. Chromatin sites undergoing active transcription are associated with histones exhibiting specific post-translational modifications, including trimethylation of the fourth lysine of histone H3 (H3K4me3) (Berr et al. 2010). The presence of H3K4me3 was initially found to correlate with an inability to establish *de novo* DNA methylation (Liu et al. 2021; Gallego-Bartolomé et al. 2019). It is now known that H3K4me3 actively recruits DNA demethylases to associated genomic loci (Wang et al. 2025). Given the differential levels of *de novo* methylation observed at the *MeSWEET10a* and *nCBP* promoters, we hypothesized that differential abundance of H3K4me3 at the respective sites might correlate to differential targeted methylation. The three promoters in the triple-targeted line 117 and wild-type cassava were evaluated for H3 and H3K4me3 association by ChIP-seq using anti-H3 and anti-H3K4me3 antibodies, respectively. As expected, H3K4me3 signal was notably abundant at the 5’end of both *nCBP* genes (Figure 4d) while no enrichment was observed at the *MeSWEET10a* locus, while H3 signal was consistent across all three regions.

### Initial assessment of inheritance and stability of *de novo* methylation at *MeSWEET10a*

To assess heritability of *de novo* methylation established using either the DRMcd or MQ1v-based tools after one meiotic generation, T0 lines with single insertions were identified and planted at a field site in Hilo, HI for self-crossing. Over 100 F1 individuals from five independent lines were evaluated for the presence of the methylation-inducing transgene and DNA methylation at the *MeSWEET10a* locus (Table 1). Seeds were collected and germinated from four different lines with the dCas9-DRMcd technology and one line with the dCas9-MQ1v technology. In total, 63 F1 progeny were positive and 48 were negative for the transgene, representing a ratio of ∼5:4 versus the expected ratio of 3:1. As expected, all 63 F1 individuals with the transgene had detectable methylation at *MeSWEET10a* (Figure 5a). Of the 49 transgene-free individuals, 41 had no detectable methylation, while 7 had detectable, albeit lower, levels of methylation compared to the parental lines or the transgene-positive F1 plants (Figure 5a). Only CpG methylation was observed in the transgene-negative F1 progeny. We selected two transgene-negative F1 progeny, P3881 and P3918, and monitored methylation at the *MeSWEET10a* promoter over time and observed that in both cases, methylation diminished and was no longer detectable by 15 months after germination (Figure 5b). Taken together, the data presented here demonstrate that Cas9-based tools can target de novo genome methylation. However, establishing methylation at a target locus is not sufficient to generate heritable epialleles. Additional research is required to uncover the molecular requirement for heritability of novel epialleles.

**Figure 5:**
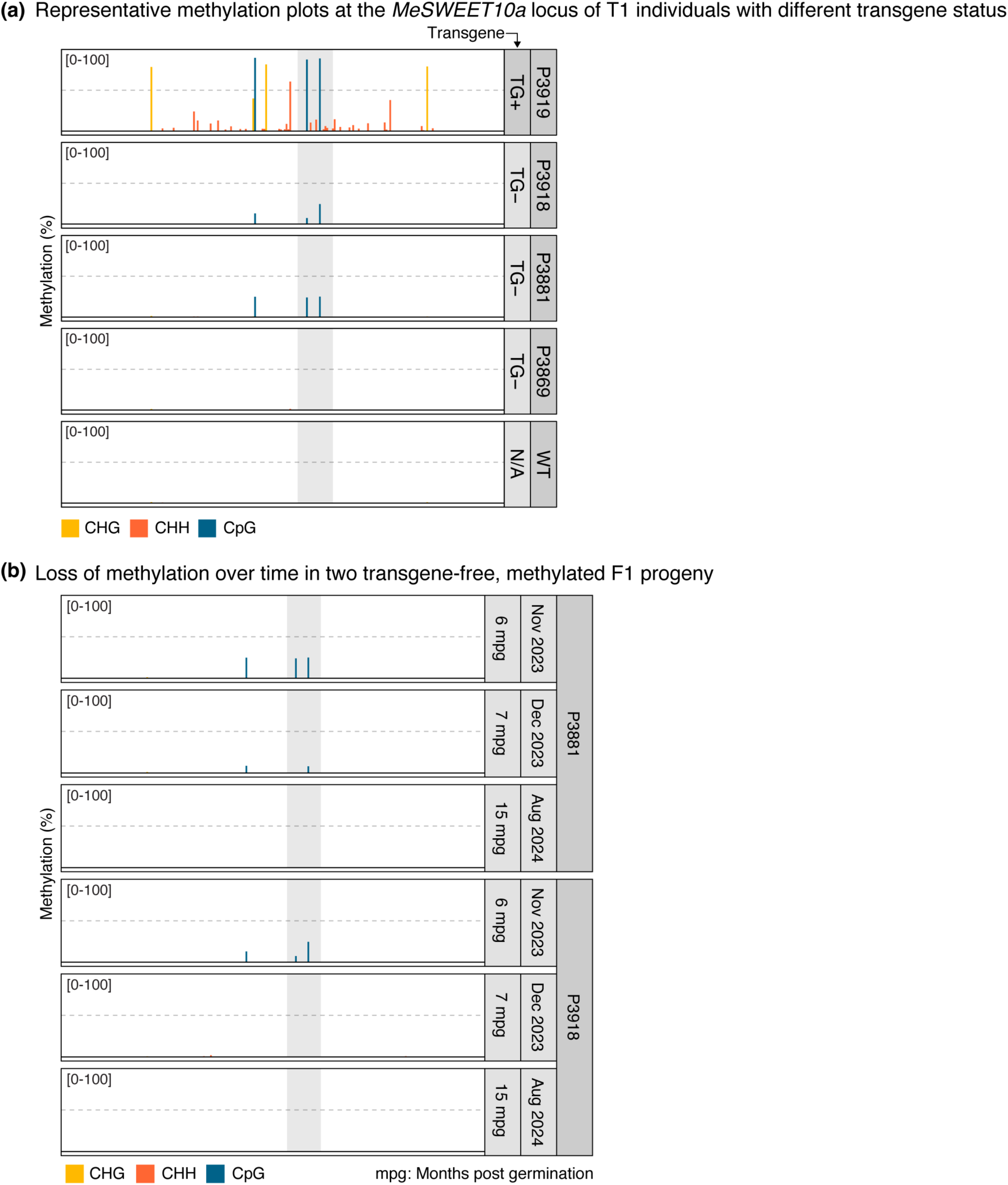
Methylation signal at the *MeSWEET10a* promoter in select T1 individuals. (a) Representative methylation profiles of T1 progeny that are (P3919) transgene positive, (P3918 and P3881) transgene negative and methylation positive, and (P3869) transgene negative and methylation negative; wildtype is an untransformed control. (b) Methylation signal in two T1 progeny, P3881 and P3918, measured over 15 months post germination (mpg).

**Table 1:**
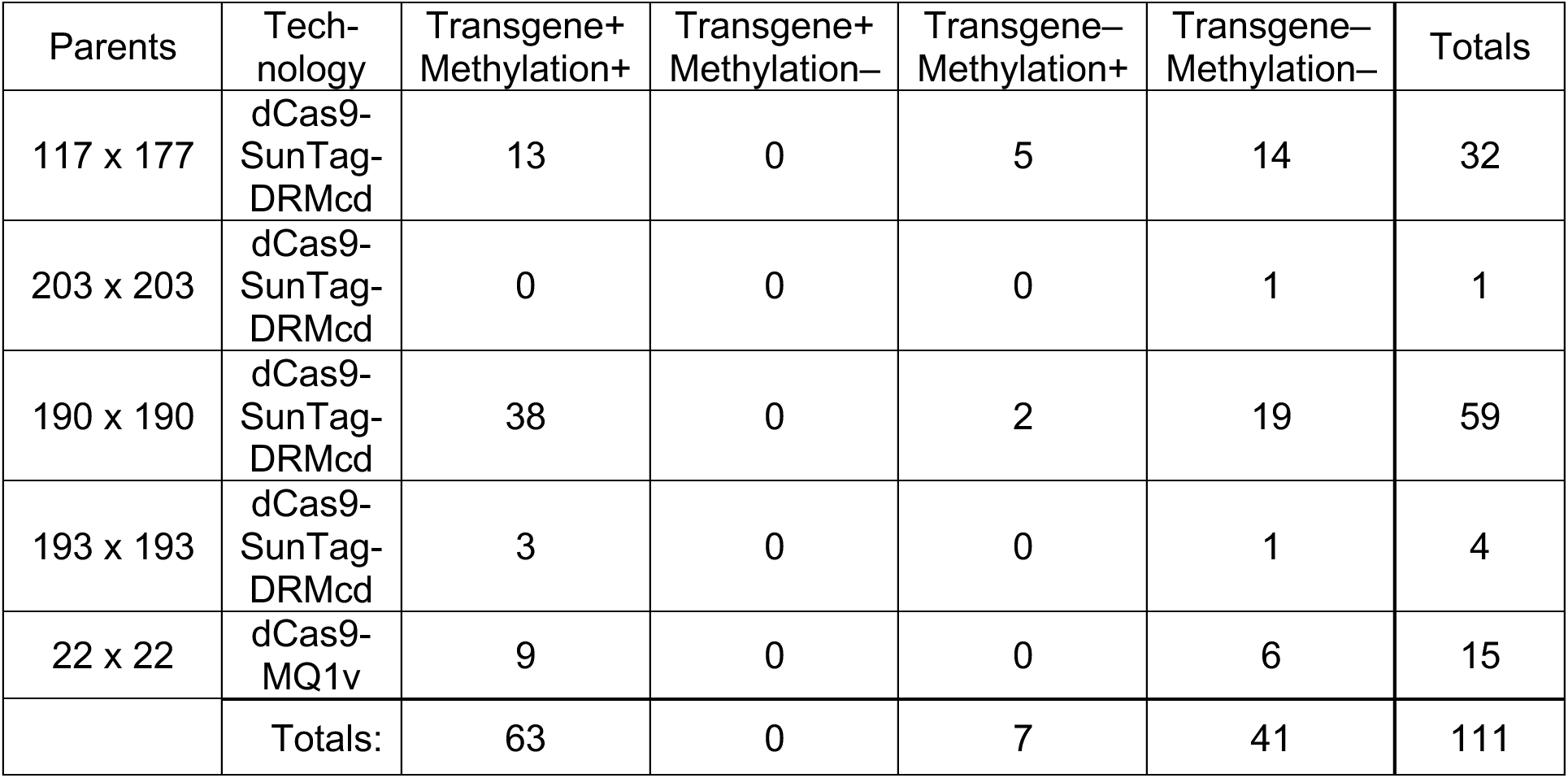
Summary transgene and methylation profiles for F1 progeny from self-crosses of *de novo* methylated T0 lines.

## Discussion

Previously, we demonstrated that directing DMS3 to the *MeSWEET10a* promoter resulted in robust methylation of the TAL20 effector-binding element (EBE), thereby preventing TAL20-mediated activation of *MeSWEET10a* expression and subsequent CBB-induced water-soaking (Veley et al. 2023). DMS3 mediates DNA methylation by recruiting the RdDM machinery for *de novo* methylation and resulted in near complete methylation at two CG positions within the TAL20 EBE and one position 22 bp upstream of the EBE. In the current study, two additional mechanisms (DRMcd and MQ1v) were tested in cassava to direct DNA methylation to the *MeSWEET10a* promoter, to analyze effects on susceptibility to CBB, and to assess heritability of targeted methylation in the absence of the targeting factors. Additionally, we also tested the simultaneous targeting of three genetic loci, *MeSWEET10a*, *nCBP-1* and *nCBP-2*, by the dCas9-SunTag-DRMcd system. Using DRMcd to target promoter sites resulted in methylation in all three cytosine contexts at all three loci, but with strong CG methylation occurring only at the *MeSWEET10a* promoter. Targeting MQ1v, a CG-specific methyltransferase, to *MeSWEET10a* resulted in strong methylation at only CG positions in and around the target site. As seen with ZF-DMS3, both MQ1v and DRMcd-mediated methylation of *MeSWEET10a* attenuated both the induction of *MeSWEET10a* by TAL20 and the water soaking symptom phenotype during CBB infection. Collectively with prior published results, we conclude that DNA methylation of promoter regions can be guided and catalyzed in cassava using several different tools, and that phenotypes consistent with repression of target genes can be obtained.

This study also revealed differences in the extent of targeted DNA methylation at different loci. In triple locus-targeted plants, the degree of overall methylation, and CG methylation particularly, was consistently higher at *MeSWEET10a* compared to both *nCBP-1* and *nCBP-2*. In ChIP-seq experiments, chromatin at the *nCBP-1* and *nCBP-2* target sites appeared to have stronger association with methylation-antagonistic H3K4me3 than did chromatin at the targeted *MeSWEET10a* site. Using Arabidopsis, it was shown that delivery of the CG-methyltransferase MQ1v with TRBIP1, a histone editor that facilitates H3K4me3 removal, results in an increase in *de novo* methylation and ultimately silencing of the *fwa* locus (Wang et al. 2025). Deployment of TRBP1 in cassava, either via a SunTag system construct or a direct fusion to dCas9, could be attempted to improve the deposition of *de novo* methylation, particularly at active promoters with H3K4me3 marks. Additionally, identifying strong promoters that enhance expression of epigenome editors in target plant species may also improve epigenome editing efficiency (Gardiner et al. 2020). These modifications may be important for maximizing efficiency, stability, and heritability of epigenome edits, and helpful in making targeted epigenome editing a viable approach to crop improvement.

We also evaluated heritability of *de novo* DNA methylation mediated by the DNA methyltransferases DRM and MQ1v after one meiotic generation (T1). As expected, methylation profiles remained robust in T1 plants co-segregating with methylation-inducing transgenes. Among all T1 plants lacking the transgene, only 14.6% had detectable DNA methylation at the target sites, and in two of those plants tested, methylation was lost over 17 months of vegetative growth. Interpretation of these results should be considered with caution, however, as others reported that DNA methylation can be enhanced over successive meiotic generations (Bond and Baulcombe 2015; Teixeira et al. 2009). Indeed, inheritance of methylation targeted to the *fwa* epiallele was not observed in a null segregant in Arabidopsis until the T2 generation with the dCas9-SunTag-DRMcd tool and the T3 generation with the dCas9-MQ1v tool (Papikian et al. 2019; Ghoshal et al. 2021). As such, passage of transgenic T1 cassava individuals through one or more rounds of self-fertilization may similarly reinforce *de novo* methylation and inheritance, ultimately resulting in stable maintenance of methylation. To be fully explored, these experiments will require long-term commitment of five to ten years given the long generation and propagation times of cassava.

In summary, we have demonstrated that two different CRISPR-based systems, using two different methyltransferases, can successfully target *de novo* methylation to the promoter region of the CBB susceptibility gene *MeSWEET10a*, resulting in attenuated symptoms. Specifically, robust CpG methylation was observed at this locus in the presence of either DRMcd or MQ1v. We also demonstrated targeting of DRMcd to multiple genetic loci and observed an increase in methylation at both the *nCBP-1* and *nCBP-2* promoters, albeit a reduced CpG signal than observed at *MeSWEET10a*. The lower methylation levels at the *nCBP*s correlated with the presence of H3K4me3 signal, a known antagonist of methylation, and thus we propose co-deployment of methylation enzymes with known H3K4me3-demethylases to improve deposition of *de novo* methylation. Inheritance of epigenetic marks in the absence of transgenic reagents is the ideal product. Here we observed methylation at the *MeSWEET10a* promoter in transgene-free T1 individuals that diminished over time; however, it remains an open question whether additional generations could result in maintenance of stable methylation at the promoters of these important S genes.

## Methods

### Construct design and cloning

A pair of gRNAs, gRNA4 and gRNA5, were designed to target the *MeSWEET10a* promoter (Table 2) and cloned into the KpnI site of the dCas9-SunTag/scFvSunTag-DRMcd epigenome editing binary vector from (Papikian et al. 2019) to generate the DRMcd-based constructs in Lines 190, 193, and 203. The ΔDRM-based construct contained the same gRNAs but lacked the DRMcd enzyme. To target the *nCBP-1* and *nCBP-2* promoters, a pair of gRNAs for each locus used previously (Z.-J. D. Lin et al. 2025) were cloned downstream of the *MeSWEET10a* gRNAs. For the MQ1-based technologies, the same two *MeSWEET10a* gRNAs from the DRM-based constructs were used; the MQ1v and dMQ1 constructs were from (Ghoshal et al. 2021).

**Table 2:**
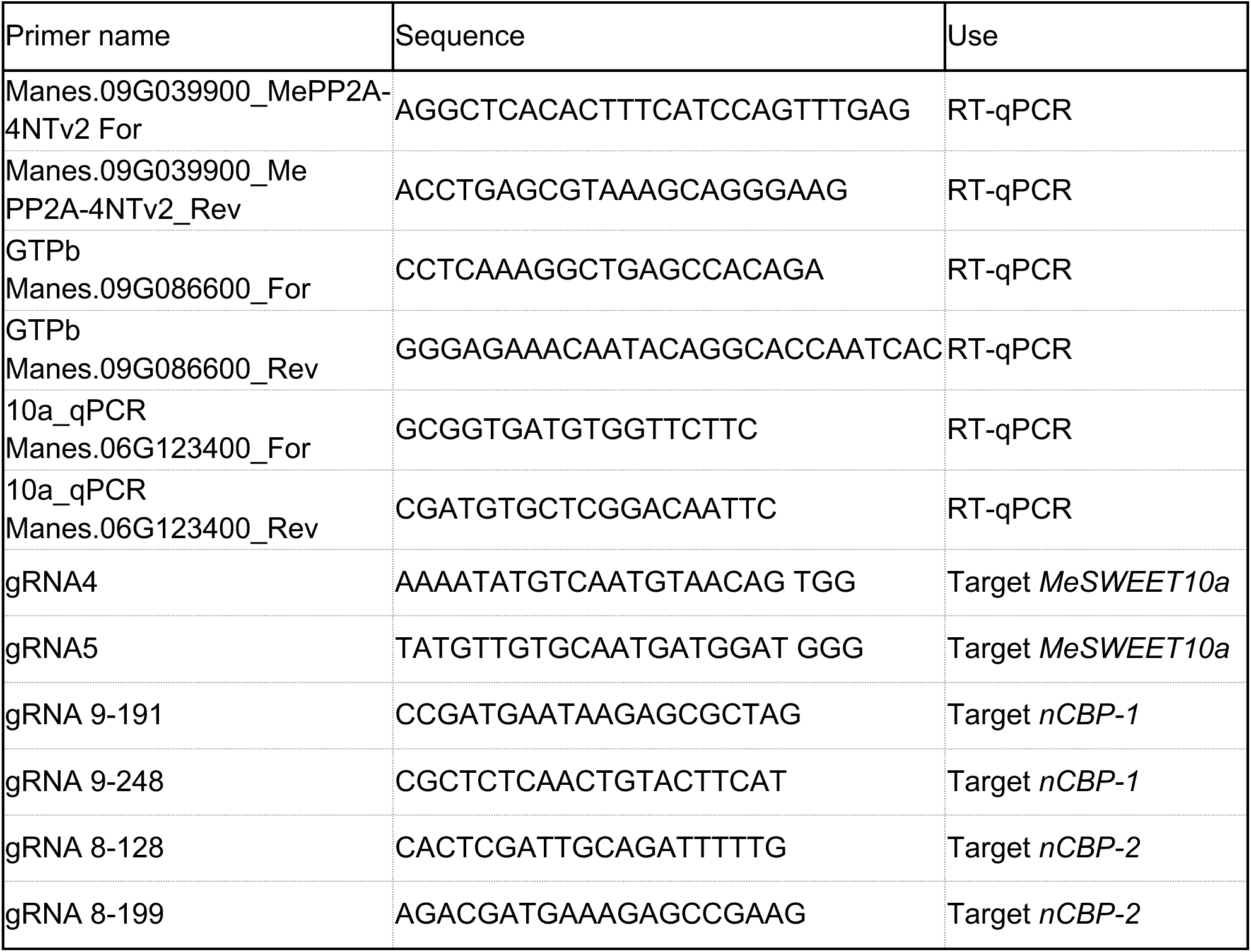
Primers and gRNA sequences.

### Production of plants and growth conditions

Transgenic cassava lines expressing either the DRM-based technologies or MQ1-based technologies were generated in the cultivar TME419 as previously described (Taylor et al. 2012; Veley et al. 2023).

### PCR bisulfite sequencing library preparation and analysis

To evaluate *de novo* methylation at target loci, PCR-based BS-seq (ampBS-seq) was used as described previously with read counts of methylated cytosines were generated using the epigenomic analysis plug-in in CLC Genomics Workbench. (Ghoshal et al. 2021; Veley et al. 2023; Z.-J. D. Lin et al. 2025). Primers used to amplify the *MeSWEET10a*, *nCBP-1*, and *nCBP-2* promoters are in Table 2.

### *Xanthamonas* infection assays

Wildtype and ΔTAL20 *Xanthamonas* strains were grown and inoculated onto cassava lines as previously described (Veley et al. 2023). Plants were sampled at 48 hours post-inoculation for RNA and DNA extraction for RT-qPCR and ampBS-seq respectively or at 96 hours post-inoculation for image-based analyses. Total RNA was extracted using the Spectrum™ Plant Total RNA Kit (Millipose-Sigma) as described previously (Veley et al. 2023; Z.-J. D. Lin et al. 2025) and primers used for RT-qPCR are in Table 2. Similarly, image-based analysis of water-soaking area size and color intensity from inoculated cassava plants was done as described previously (Veley et al. 2023).

### H3K4me3 ChIP-sequencing

ChIP-Seq was performed as previously described (Wang et al. 2025) with the following modifications. A total of 5 grams of tissue was collected from the first two fully formed leaves of approximately 6 month old greenhouse grown cassava plants. Ground tissue was resuspended in Nuclear Isolation Buffer and incubated at 4°C for 25 minutes while rotating and the reaction was terminated with fresh glycine. After filtering the samples and the centrifugation step, the pellet was resuspended in 2mL of Lysis Buffer 1 (50 mM Tris-HCl pH=8.0, 10 mM EDTA, 1.0% SDS, 0.1 mM PMSF, 5 mM benzamidine, 1x cOmplete Mini EDTA- free Protease Inhibitor Cocktail). Following an incubation at 4°C for 20 minutes, centrifugation for 10 minutes, the pellet was then resuspended in 2mL of Lysis Buffer 2 (50mM Tris-HCl pH= 8.0, 10 mM EDTA, 0.1% SDS, 0.1 mM PMSF, 5 mM benzamidine, 1x cOmplete Mini EDTA- free Protease Inhibitor Cocktail). Samples were sheared using the Covaris S220 Biorupter at 150 watts, 20% duty factor, 200 cycles/bursts, and 600 seconds in a 2 ml Covaris Tube (catalog number 520056, Covaris) in an adaptor (catalog number 500199, Covaris). The lysate was centrifuged at 4°C for 10 minutes and the supernatant moved to new tube and centrifuged for 10 minutes at 4°C. The supernatant was then incubated with 2 mL of ChIP Dilution Buffer (1.1% Triton X-100, 1.2 mM EDTA, 16.7 mM Tris-HCl pH=8.0, 167 mM NaCl, 0.1 mM PMSF, 5 mM benzamidine, 1 Protease Inhibitor Cocktail) after which the sample was divided into 4 separate aliquots: 100 ul for the input sample and the remaining 3.9 mL was split into 3 tubes for either bead only control, or 1 µl of anti-H3K4me3 (catalog number 04-745R, Millipore), or 1 µl of anti-H3 antibody (catalog number ab1791, abcam). After overnight incubation the DNA was precipitated and the pellet is washed with 70% ethanol, and resuspended in13 ul of Buffer EB (catalog number 19086, Qiagen). Libraries were constructed with the NuGen Ultralow V2 kit and sequenced on an Illumina NovaSeq X plus sequencer.

### Bioinformatic analysis of H3K4me3 data

Raw sequencing data from the ChIP-seq experiment was trimmed using trim_galore (https://github.com/felixkrueger/trimgalore). Trimmed reads were then aligned to the haplotype-resolved cassava genome of variety TME204 (Qi et al. 2022) using bowtie2 (version 2.5.3; (Langmead and Salzberg 2012) and samtools (version 1.19.2; (Danecek et al. 2021)) was used to convert to bam format. After using picard MarkDuplicates (version 3.1.1; http://broadinstitute.github.io/picard), deeptools (version 3.5.5; (Ramírez et al. 2016)) was used to normalize the data using RPKM and generate bedfiles of the regions of interest.

## Data availability and code availability

Raw ChIP-seq sequencing data generated during this study is available at GEO (Accession number: GSE329762). Raw images of Xpm inoculated leaves, processed data tables, and R code used for generation of all figures is available at FigShare (https://figshare.com/s/3511a3f7bcd4c21ea70c).

## Generation of T1 progeny

Plantlets in tissue culture were sent to facilities in Hilo, Hawaii for hardening off and planting at a field site, as previously described (Elliott et al. 2024). When flowering occurred, flowers were protected from open pollination, and instead specific females and males were selected for crossing (or selfing). Seeds that resulted from these hand pollinations were collected 2-3 months later, along with their identifying tag. After drying down, they were packaged and sent to the Donald Danforth Plant Science Center in Saint Louis, MO for germination and analysis of methylation and transgene status.

## Acknowledgements

We are grateful for the scientific advice from Drs. Nigel Taylor and Williams Esuma. We also acknowledge Dr. Noah Fahlgren, Director of the Data Science lab at DDPSC, and the additional members of our labs who contributed to this work in various ways, including Joshua Sumner, Drs. Xinggou Zheng and Kiona Elliott. This work was funded by the Bill and Melinda Gates Foundation (OPP1210659, R.B.S., S.E.J., S.M.W., J.C.C.). We dedicate this work to the memory of Lukas Kambic and honor his enthusiasm for all things plant science.

## Author contributions

Conceptualization: R.S.B, S.E.J.,S.M.W., J.C.C.,Z.D.L,K.M.V. Data curation: K.B.G., Z.D.L,K.M.V,M.Y. Formal analysis: K.B.G., Z.D.L,K.M.V,M.Y. Funding acquisition: R.S.B, S.E.J.,S.M.W., J.C.C. Investigation: K.B.G., Z.D.L., K.M.V., M.K.S., M.Y., K.Z., G.J., E.M., G.L.H.,S.F.,Y.H. Methodology: Z.D.L.,K.M.V,M.K.S.,G.L.H. Project administration: K.B.G., R.S.B, S.E.J.,S.M.W., J.C.C. Resources: K.B.G.,Z.D.L.,K.M.V.,M.K.S.,G.L.H., R.S.B, S.E.J., S.M.W., J.C.C. Software: K.B.G. Supervision: K.B.G.,Z.D.L.,K.M.V., M.Y.,R.S.B, S.E.J., S.M.W., J.C.C. Validation: Z.D.L.,K.M.V.,G.J.,E.M. Visualization: K.B.G. Writing – original draft: K.B.G., Z.D.L.,M.K.S. Writing – review & editing: K.B.G.,Z.D.L.,K.M.V., R.S.B, S.E.J., S.M.W., J.C.C.

---

**Supplemental Figure 1:**
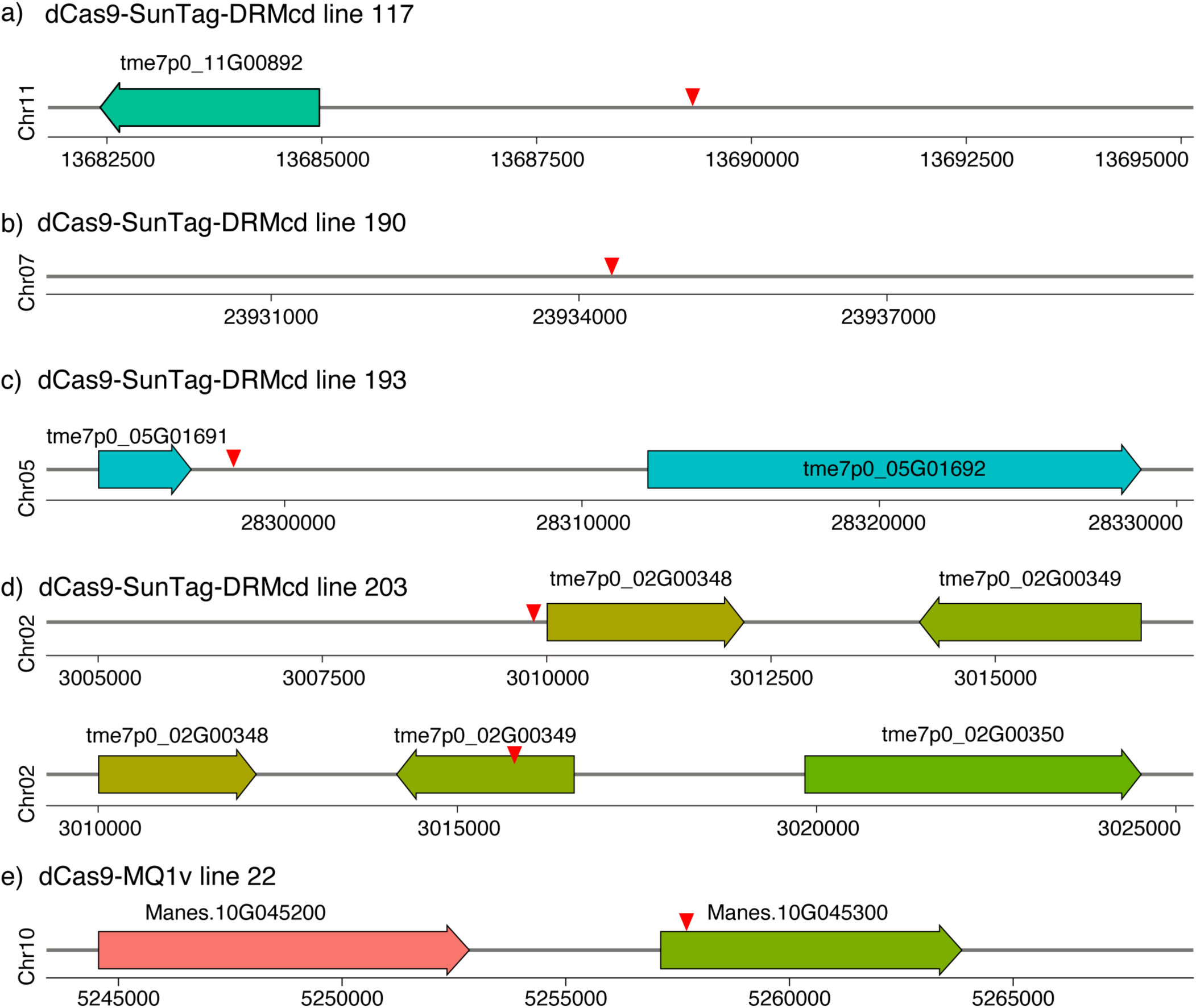
Transgene insertion location and number for dCas9-SunTag-DRMcd and dCas9-MQ1v lines. (a-d) Location of transgene insertions relative to the TME7 genome (Mansfeld et al., 2021) and (e) location relative to the v8 *Manihot esculenta* genome at Phytozome. (a) Location of transgene in dCas9-SunTag-DRMcd Line 117; (b) location of transgene in dCas9-SunTag-DRMcd Line 190; (c) location of transgene in dCas9-SunTag-DRMcd Line 193; (d) location of 2 transgene insertion in dCas9-SunTag-DRMcd Line 203, only 5,920 bp apart; (e) location of transgene in dCas9-MQ1v Line 22.

**Supplemental Figure 2:**
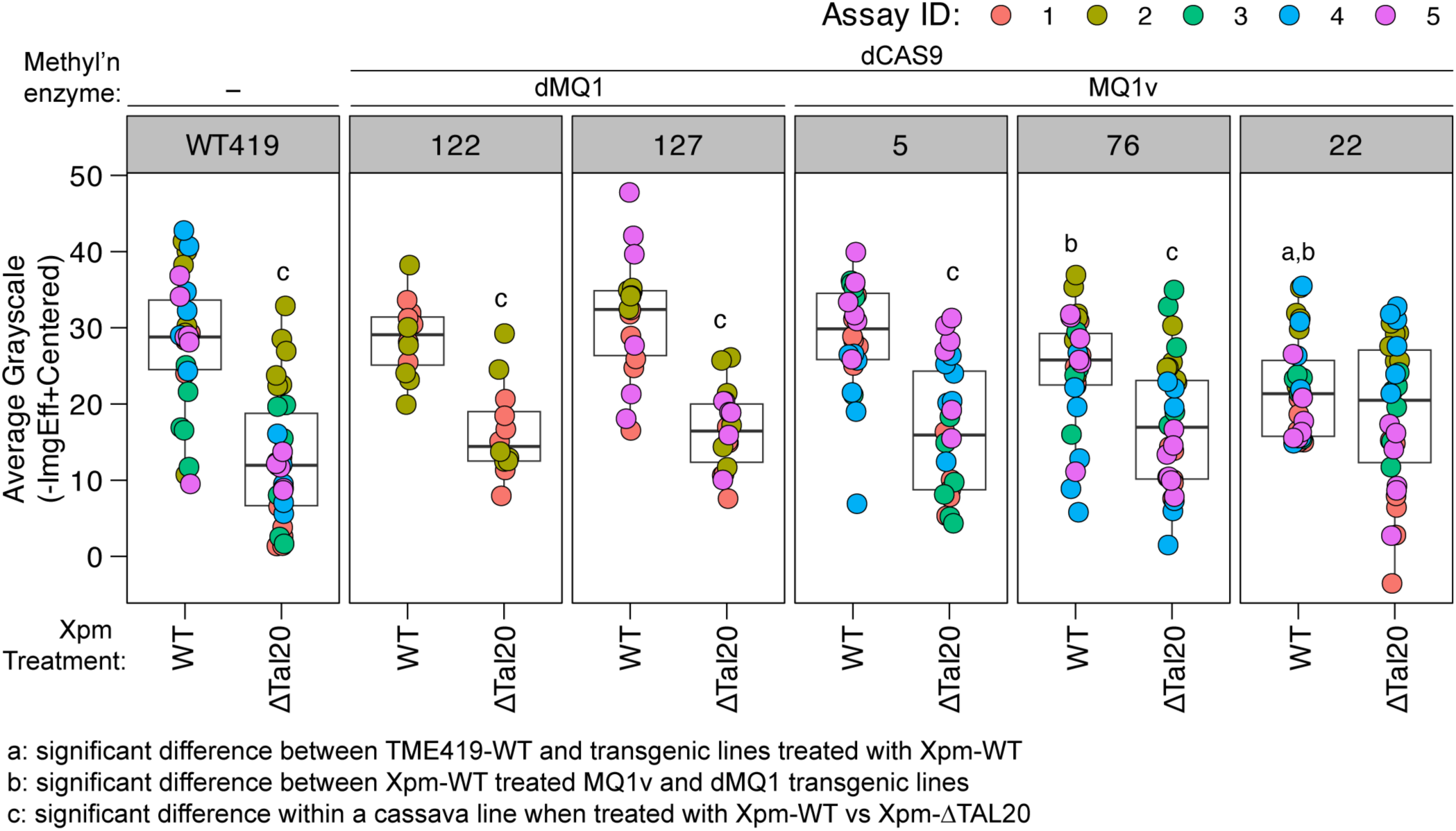
dCas9-MQ1v lines with targeted CG methylation to the cassava *MeSWEET10a* promoter have attenuated CBB symptoms. Intensity of water-soaking lesions from five assays; values plotted are the negative mean gray-scale value for the water-soaked region relative to the average of the mock-treated samples. Calculated *p* values from two-sided Kolmogorov–Smirnov test are shown indicated above the boxplots and are defined at the bottom of the figure.

**Supplemental Figure 3:**
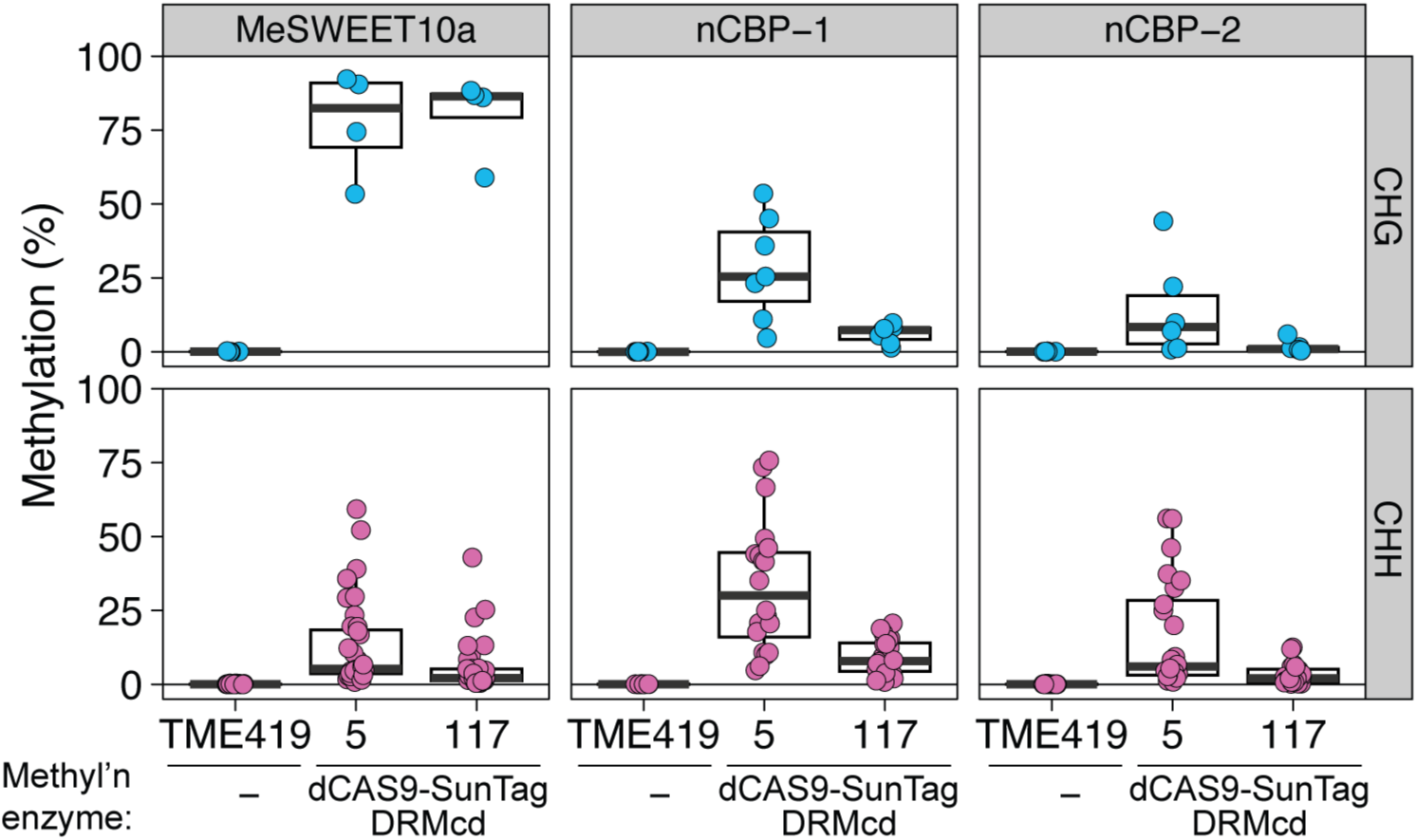
CHG and CHH methylation signal at three simultaneously targeted loci. Methylation at the *MeSWEET10a*, *nCBP-1*, and *nCBP-2* promoter regions in the two other cytosine contexts (CGH, CHH) as measured by PCR-based bisulfite sequencing (ampBS-seq) in two independent transgenic lines expressing dCas9-SunTag-DRMcd (5 and 117) and wildtype TME419. Y-axis is percent methylated from 0 to 100.

